# DNA damage causes ATM-dependent heterochromatin loss leading to nuclear softening, blebbing, and rupture

**DOI:** 10.1101/2024.05.24.595790

**Authors:** Nebiyat Eskndir, Manseeb Hossain, Marilena L Currey, Mai Pho, Yasmin Berrada, Andrew D Stephens

**Affiliations:** Biology Department, University of Massachusetts Amherst, Amherst, MA; Molecular and Cellular Biology, University of Massachusetts Amherst, Amherst, MA 01003, USA

## Abstract

The nucleus must maintain stiffness to protect the shape and integrity of the nucleus to ensure proper function. Defects in nuclear stiffness caused from chromatin and lamin perturbations produce abnormal nuclear shapes common in aging, heart disease, and cancer. Loss of nuclear shape via protrusions called blebs leads to nuclear rupture that is well-established to cause nuclear dysfunction, including DNA damage. However, it remains unknown how increased DNA damage affects nuclear stiffness, shape, and ruptures, which could create a negative feedback loop. To determine if increased DNA damage alters nuclear physical properties, we treated MEF cells with DNA damage drugs cisplatin and bleomycin. DNA damage drugs caused increased nuclear blebbing and rupture in interphase nuclei within a few hours and independent of mitosis. Micromanipulation force measurements reveal that DNA damage decreased chromatin-based nuclear mechanics but did not change lamin-based strain stiffening at long extensions relative to wild type. Immunofluorescence measurements of DNA damage treatments reveal the mechanism is an ATM-dependent decrease in heterochromatin leading to nuclear weaken, blebbing, and rupture which can be rescued upon ATM inhibition treatment. Thus, DNA damage drugs cause ATM-dependent heterochromatin loss resulting in nuclear softening, blebbing, and rupture.

## Introduction

The nucleus is the organelle that houses the genome and its essential functions. This organelle relies on both the chromatin at the interior and lamins at the nuclear periphery to maintain its mechanical stiffness which determines nuclear shape and integrity. Loss of nuclear stiffness, shape, and integrity occurs in many human afflictions including aging, muscular dystrophy, heart disease, and many cancers (Stephens *et al*., 2019a; Kalukula *et al*., 2022). The last decade has revealed that loss of stiffness, shape, and integrity leads to nuclear dysfunction. Thus, nuclear physical properties are not just a hallmark of human disease but a major cause of its dysfunction. Specifically, nuclear blebbing and rupture cause increased DNA damage (Denais *et al*., 2016; Irianto *et al*., 2016; Raab *et al*., 2016; Chen *et al*., 2018; Xia *et al*., 2018; Stephens *et al*., 2019b; Earle *et al*., 2020; Nader *et al*., 2021; Shah *et al*., 2021; Pho *et al*., 2023), changes in transcription (De Vos *et al*., 2011; Helfand *et al*., 2012; Berg *et al*., 2023), and perturbations to cell cycle control (Pfeifer *et al*., 2018). Increased DNA damage is a consistently reported outcome of loss of nuclear integrity via rupture resulting from loss of nuclear stiffness and shape. However, it remains to be determined if nuclear dysfunctions such as DNA damage can induce loss of nuclear stiffness, shape, and integrity causing a negative feedback loop to drive disease severity.

Many recent studies have investigated the effect of increased DNA damage on nuclear stiffness, shape, and integrity. These studies imply that both major DNA damage repair kinases affect nuclear physical properties. Ataxia-telangiectasia mutated (ATM) repairs DNA strand breaks and ATM- and Rad3-Related (ATR) responds to broad DNA damage (Maréchal and Zou, 2013). It was first reported that increased DNA damage resulted in decreased nuclear stiffness due to chromatin reorganization (Dos Santos *et al*., 2021). Though the mechanism of action and the consequences remain unclear. One insight was the ATM inhibition reversed the loss of the nuclear stiffness (Dos Santos *et al*., 2021). Another report states that ATM inhibition decreases lamin levels and thus decreases lamin-based nuclear stiffness (Shah *et al*., 2022). Thus, the field disagrees, wherein one report that shows DNA damage with ATM inhibition rescues (increases) nuclear stiffness while another report shows non-DNA damage induced ATM inhibition makes the nucleus weaker. Finally, another report states that ATR activation by DNA damage results in phosphorylation of lamin A/C to cause rupture (Kovacs *et al*., 2023) while another reports depletion of ATR disrupts cell mechanics and migration (Kidiyoor *et al*., 2020). In short, many of the most influential labs in nuclear mechanobiology agree that DNA damage and repair signaling affects nuclear mechanics, shape, and integrity, but the mechanisms remain unclear.

While the field of nuclear mechanobiology works to understand this new phenomenon, the field of DNA damage has concluded that ATM has a pivotal role in both DNA damage repair and chromatin compaction state through heterochromatin (Li *et al*., 2020). Initial DNA damage recruits heterochromatin forming proteins Kap-1/HP1/Suv39h1 to deposit constitutive heterochromatin marker H3K9me^3^ (Ayrapetov *et al*., 2014) which aids activation of ATM through Tip60 (Sun *et al*., 2009). However, ATM phosphorylates Kap-1 to dissociate it and the heterochromatin machinery (Ziv *et al*., 2006; Ayrapetov *et al*., 2014) to aid chromatin relaxation. Separately, at DNA damage sites PARP1 recruits the histone demethylases KDM4B/D that remove H3K9me^3^ methyl groups to decreased heterochromatin and decompact the underlying chromatin for proper DNA damage repair (Young *et al*., 2013; Khoury-Haddad *et al*., 2014). This is important because heterochromatin has been shown to be an essential nuclear mechanical component that controls nuclear shape and integrity (Stephens *et al*., 2017, 2018, 2019b; Nava *et al*., 2020; Williams *et al*., 2020; Strom *et al*., 2021). Thus, DNA damage repair via ATM has a dedicated pathway to decompact heterochromatin that could alter nuclear physical properties.

To determine the effect of increased DNA damage on nuclear shape, integrity, and stiffness, we induced DNA damage via drugs and measured these key physical properties of nuclei. Mouse embryonic fibroblast (MEF) cells were treated with DNA damage inducing drugs cisplatin to cause DNA adducts and bleomycin to cause DNA strand breaks. We then measured nuclear blebbing and ruptures via time lapse imaging, which increased in all DNA damage drug treatments. Next, we used micromanipulation force measurements to separate the nuclear stiffness contributed by chromatin at short extensions and lamins at long extension via strain stiffening (Stephens *et al*., 2017; Hobson *et al*., 2020; Currey *et al*., 2022). To determine the specific nuclear component modulated by DNA damage, we measured levels of histone modification states and lamins. Finally, we determined the mechanism by inhibiting ATM, a major kinase responsible for DNA damage response, which resulted in a rescue of nuclear shape and stability. Our studies show support for previous chromatin-based stiffness findings (Dos Santos *et al*., 2021) while providing novel mechanism where ATM-dependent decreased heterochromatin is specifically responsible for loss of nuclear mechanics, shape, and integrity. Thus, we provide a clear negative feedback loop between DNA damage and nuclear rupture which causes more DNA damage.

## Results and Discussion

### Treatment with DNA damage inducing drugs causes nuclear blebbing and rupture

To induce varying levels and types of DNA damage, we treated mouse embryonic fibroblast (MEF) cells with DNA damaging agents. Using different DNA damage treatments allows us to assess the importance of different types of DNA damage. Specifically, we treated with cisplatin at low and high (5 vs. 25 µM) levels to cause DNA adducts (Basu and Krishnamurthy, 2010) and a radiomimetic bleomycin at 13 µM to cause DNA breaks (Chen and Stubbe, 2005). To measure DNA damage levels caused by each treatment, we measured the levels of DNA damage foci labeled by γH2AX immunofluorescence. Treatment for 16 hours with either low or high cisplatin (CSP) or bleomycin (BLE) showed a significant increase in γH2AX DNA damage foci as expected (**Figure 1, A and B**). Thus, we can modulate both levels and types of DNA damage.

**Figure 1.**
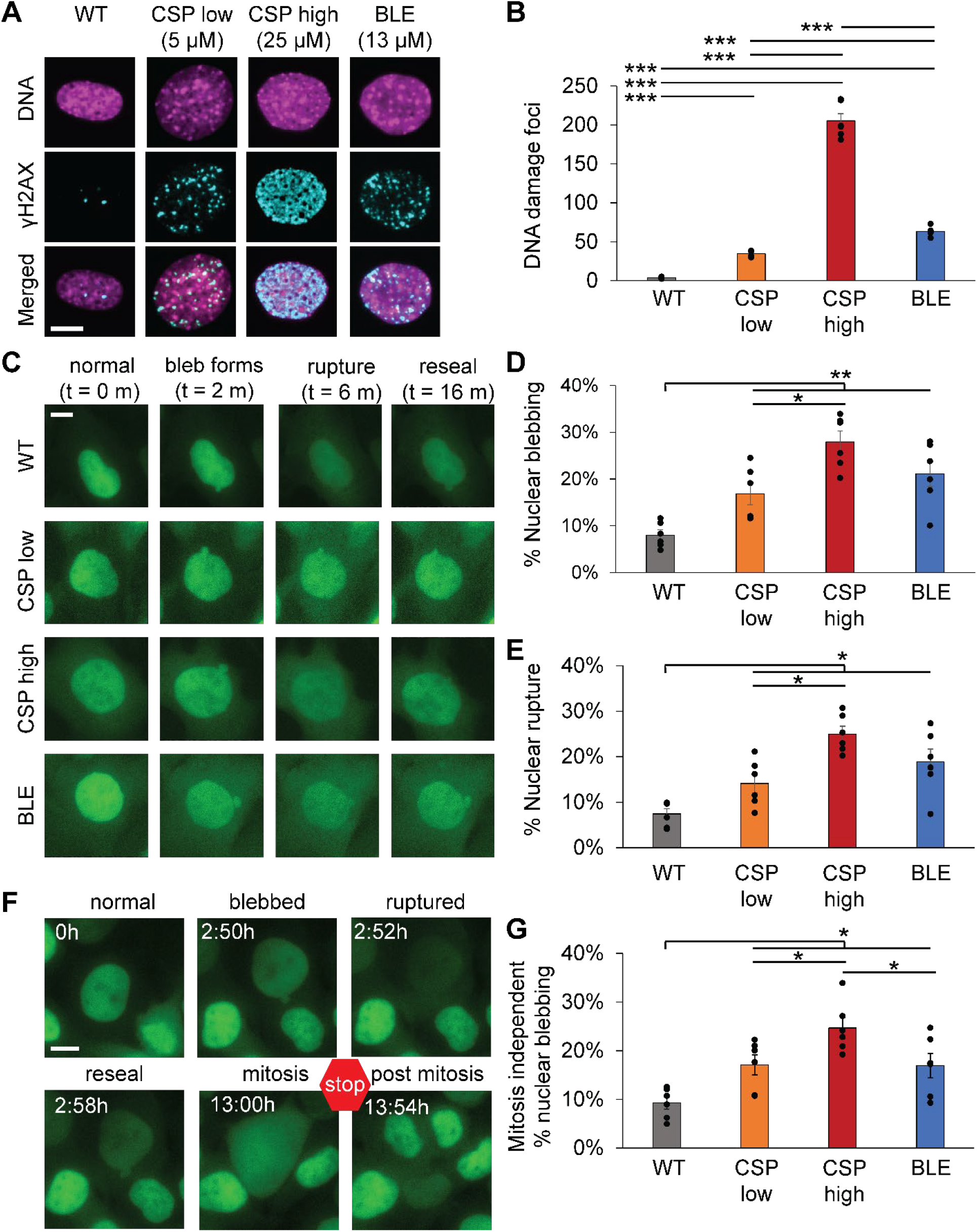
DNA damage induction causes increased nuclear blebbing and ruptures. (A) Example images of MEF nuclei DNA stained by Hoechst (magenta) and immunofluorescence for DNA damage marker gamma H2AX (cyan) for wild type (WT) and DNA damage treatment for 16 hours via cisplatin (CSP) and bleomycin (BLE). (B) Graph of quantified number of DNA damage foci for each condition with six replicates with an average n = 60, 28, 23, 51 nuclei each respectively. (C) Example images of MEF NLS-GFP from timelapse during no treatment (wild type, WT) and treatment with a DNA damage drug (CSP or BLE) for 16 hours. Images show normal nucleus into bleb formation, rupture, and resealing of the nuclear envelope with bleb remaining over 2-minute intervals throughout time lapse imaging. (D and E) Graphs of (D) nuclear blebbing and (E) nuclear rupture for wild type (WT) and DNA damage treatment for 16 hours via cisplatin (CSP) and bleomycin (BLE). (F) Example images tracking time post DNA drug treatment till mitosis. (G) Graph of percentage nuclear blebbing occurring in nuclei before first mitosis for no treatment (wild type, WT) and treatment with a DNA damage drug (CSP or BLE). Cells post mitosis are excluded from this analysis. Each condition consists of 6 replicates n = 100-300 cells each. Student’s t-test p values reported as *<0.05, **<0.01, ***<0.001, or ns denotes no significance, p>0.05. Error bars represent standard error. Scale bar = 10 µm.

DNA damage is a common dysfunction in human disease that could lead to disruption of nuclear physical properties. To determine the effect of DNA damage level and type on nuclear blebbing and rupture, we treated MEF NLS-GFP stable cells with low cisplatin, high cisplatin, and bleomycin and live-cell imaged at 2-minute intervals for 16 hours. We quantified nuclear blebbing by a protrusion > 1 µm from the nuclear body, as previously established (Stephens *et al*., 2018, 2019b). Nuclear ruptures were defined as a > 25% change in the NLS-GFP nuclear/cytoplasm ratio due to NLS-GFP concentrated in the nucleus spilling into the cytoplasm, as previously established (Pho *et al*., 2023). Untreated MEF NLS-GFP nuclei presented similarly low levels of both nuclear blebbing and rupture (8±1 %, **Figure 1C-E**). However, DNA damage induction by all treatments resulted in increased levels of both nuclear blebbing and ruptures (> 15 %, **Figure 1C-E**). The formation of nuclear blebs in interphase was followed by nuclear rupture in > 90% of cases, as previously reported (Stephens *et al*., 2019b). To rule out the effects of mitosis, we tracked nuclear blebbing in interphase nuclei from treatment up until the first mitosis (**Figure 1F**). Independent of mitosis, nuclear blebs formed during the initial interphase at significantly increased percentages compared to wild type as early as two hours post-DNA damage induction (**Figure 1, F and G**). Human fibrosarcoma HT1080 cells showed a similar significant increase in nuclear blebbing upon cisplatin and bleomycin treatment, suggesting that DNA damage provides repeatable results across cell lines (**Supplemental Figure 1**). Overall, our data reveal that DNA damage causes increased nuclear blebbing and ruptures during interphase.

Our data report the novel finding that DNA damage induction increases nuclear blebbing which leads to nuclear rupture. Interestingly, relative to wild type the amount and type of DNA damage is unimportant as all treatments resulted in a significant increase. Though there are significant differences between treatments, especially cisplatin low and high, this appears be due largely to levels of DNA damage rather than type. While the finding of nuclear blebbing is novel, increased nuclear rupture associated with DNA damage has been reported by others (Kidiyoor *et al*., 2020; Kovacs *et al*., 2023). Increased DNA damage in RPE-1 TP53-/-cells increased with a broad array of DNA damaging drugs (Kovacs *et al*., 2023) but there was no report of effects on nuclear blebbing even though example images appear to have small nuclear blebs. In a different approach, ATR depletion results in rupture under less confinement (Kidiyoor *et al*., 2020). Thus, our findings provide an advancement of our understanding of the effects of DNA damage to nuclear shape and integrity via nuclear blebs, while also being supported by recent work from others.

### DNA damage softens chromatin rigidity but not lamin strain stiffening

Nuclear blebbing and rupture is well-reported to stem from loss of nuclear stiffness from either chromatin decompaction or lamin loss (Stephens *et al*., 2019a; Kalukula *et al*., 2022). Dual micropipette single isolated nucleus micromanipulation force measurements provide the unique ability to measure the separate stiffness contributions of chromatin at short extensions and lamins at long extensions via strain stiffening (**Figure 2A**)(Stephens *et al*., 2017; Currey *et al*., 2022). This is especially important since the DNA damage and repair pathway has been reported to alter nuclear mechanics through both chromatin (Dos Santos *et al*., 2021) and lamins (Shah *et al*., 2022; Kovacs *et al*., 2023).

**Figure 2.**
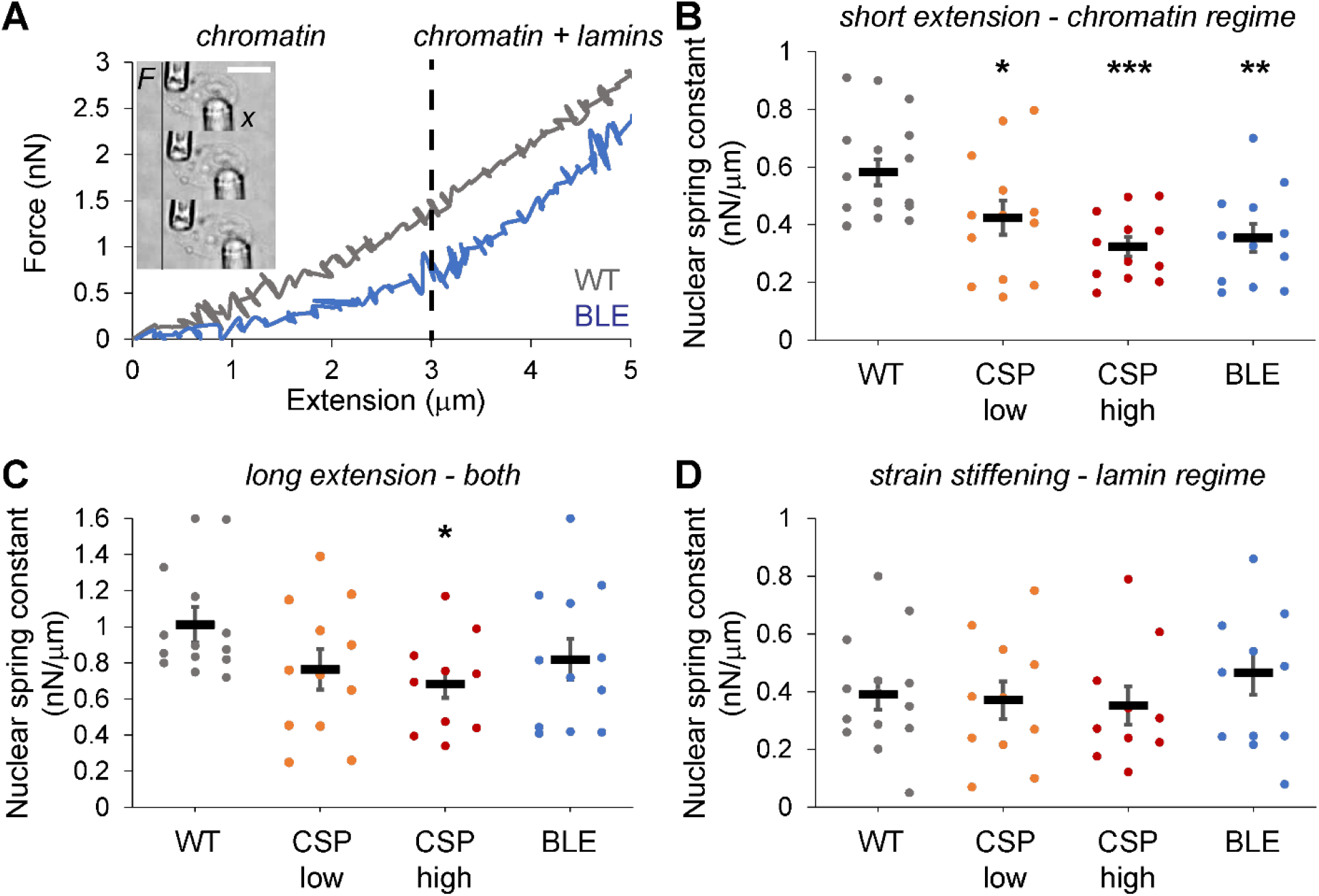
Micromanipulation force measurements reveal induced DNA damage softens chromatin rigidity but not lamin strain stiffening. (A) Example of isolated cell nucleus suspended between the force (F) pipette and the pull (x) pipette. During micromanipulation the pull pipette extends the nucleus while nucleus extension (µm) is measured by distance between the two pipettes. The force pipette’s deflection multiplied by a pre-measured bending constant provides a measure of force (nN). Example force vs. extension traces of wild type (WT, gray) and DNA damage (BLEO, blue). Micromanipulation allows for the separation of chromatin-based rigidity at short extension (< 3 µm) vs. lamin-based strain stiffening at long extension (< 3 µm). (B, C, D) Graphs of the nuclear spring constant (nN/µm) for wild type (WT) and DNA damage caused by cisplatin low (CSP 5 µM), cisplatin high (25 µM) and bleomycin (BLEO) at over different extensions. (B) Graph of nuclear spring constat for short extension (< 3 µm) chromatin-based regime (n = 18, 13, 12, 12 respectively). (B) Graph of nuclear spring constat for long extension (> 3 µm) in which both chromatin and lamins are contributing to nuclear rigidity (n = 14, 12, 10, 12 respectively). (B) Graph of nuclear spring constat for long extension strain stiffening lamin-based regime which is calculated by the difference between the short and long extension nuclear spring constant (n = 14, 12, 10, 12 respectively). Student’s t-test p values reported as *<0.05, **<0.01, ***<0.001, or ns denotes no significance, p>0.05. Error bars represent standard error. Scale bar = 10 µm.

Micromanipulation force measurements of isolated nuclei from MEF vimentin null (V-/-) cells were conducted for untreated and those treated with DNA damaging drugs. Isolated nuclei were non-specifically attached to two micropipettes where the “pull” pipette moves to extend the nucleus while the “force” pipettes deflection multiplied by a pre-measured bending constant provides a measure of force. This force in nN vs. extension in µm provides a nuclear spring constant nN/µm measured as the slope of the line (**Figure 2A**). Untreated nuclei measured a similar and consistent 0.58 ± 0.1 nN/um for the chromatin-based short extension force regime (0-3 µm extension, **Figure 2B**) with a strain stiffening increase at extensions > 3 µm, as previously reported (Stephens *et al*., 2017; Currey *et al*., 2022). DNA damage induction from all three treatments resulted in a significant decrease in the chromatin-based short extension nuclear spring constant to 0.42 – 0.32 nN/um (p < 0.05, **Figure 2B**). Long extension nuclear spring constants, which measures the mixed contribution of chromatin and lamins, trended lower for all DNA damage treatments, but only high cisplatin treatment measured a statistically significant decrease (**Figure 2C**). Next, we assayed the contribution of lamin-based strain stiffening by calculating the difference between short and long extension nuclear spring constants. Lamin-based strain stiffening spring constant showed no significant change between wild type and DNA damage treatments (**Figure 2D**). Overall, our micromanipulation force measurements provide valuable data clearly showing a significant decrease in chromatin-based nuclear rigidity upon DNA damage while lamin-based strain stiffing was unchanged.

Our novel data provides a key insight that clarifies the current literature. This data generally agrees with previous atomic force microscopy (AFM) measurements that reported a decreased nuclear rigidity upon cisplatin treatment (Dos Santos *et al*., 2021). However, our micromanipulation technique has provided the novel finding that specifically decreased chromatin rigidity is responsible for the weaker nucleus measurements. This finding is in agreement with the AFM paper which concluded DNA damage causes changes in chromatin. Furthermore, our data showing that lamin-based strain stiffening remains unperturbed does not support claims of changes in lamins, at least lamin-based nuclear mechanics, upon changes in ATM, ATR, or DNA damage (Shah *et al*., 2022; Kovacs *et al*., 2023). However, the disagreement with changes in lamins might be due largely to different approaches. Overall, many labs are finding that the DNA damage and repair pathway impacts nuclear mechanics and integrity.

### DNA damage causes ATM-dependent heterochromatin loss

Heterochromatin is a well-established mechanical element of the nucleus (Stephens *et al*., 2017) that has both been documented to change in response to DNA damage (Ziv *et al*., 2006; Ayrapetov *et al*., 2014) and lead to loss of nuclear rigidity, shape, and integrity (Stephens *et al*., 2018, 2019b). To determine if heterochromatin loss results from DNA damage treatments, we conducted immunofluorescence of H3K9me^2,3^ a constitutive heterochromatin marker. Upon DNA damage by cisplatin or bleomycin H3K9me^2,3^ heterochromatin signal significantly decreased compared to wild type (**Figure 3A-C**). Treatment of cells with both DNA damage drug and ATM inhibitor (ATMi) KU55933 resulted in a significant increase in heterochromatin relative to DNA damage treatment alone. In DNA damage treatments the euchromatin marker H3K9ac did not increase and lamin A/C levels did not significantly decrease (**Supplemental Figure 2**), suggesting they have no role in the mechanism. Taken together, this data suggests that ATM-dependent response to DNA damage causes heterochromatin loss, which can be restored upon ATM inhibition.

**Figure 3.**
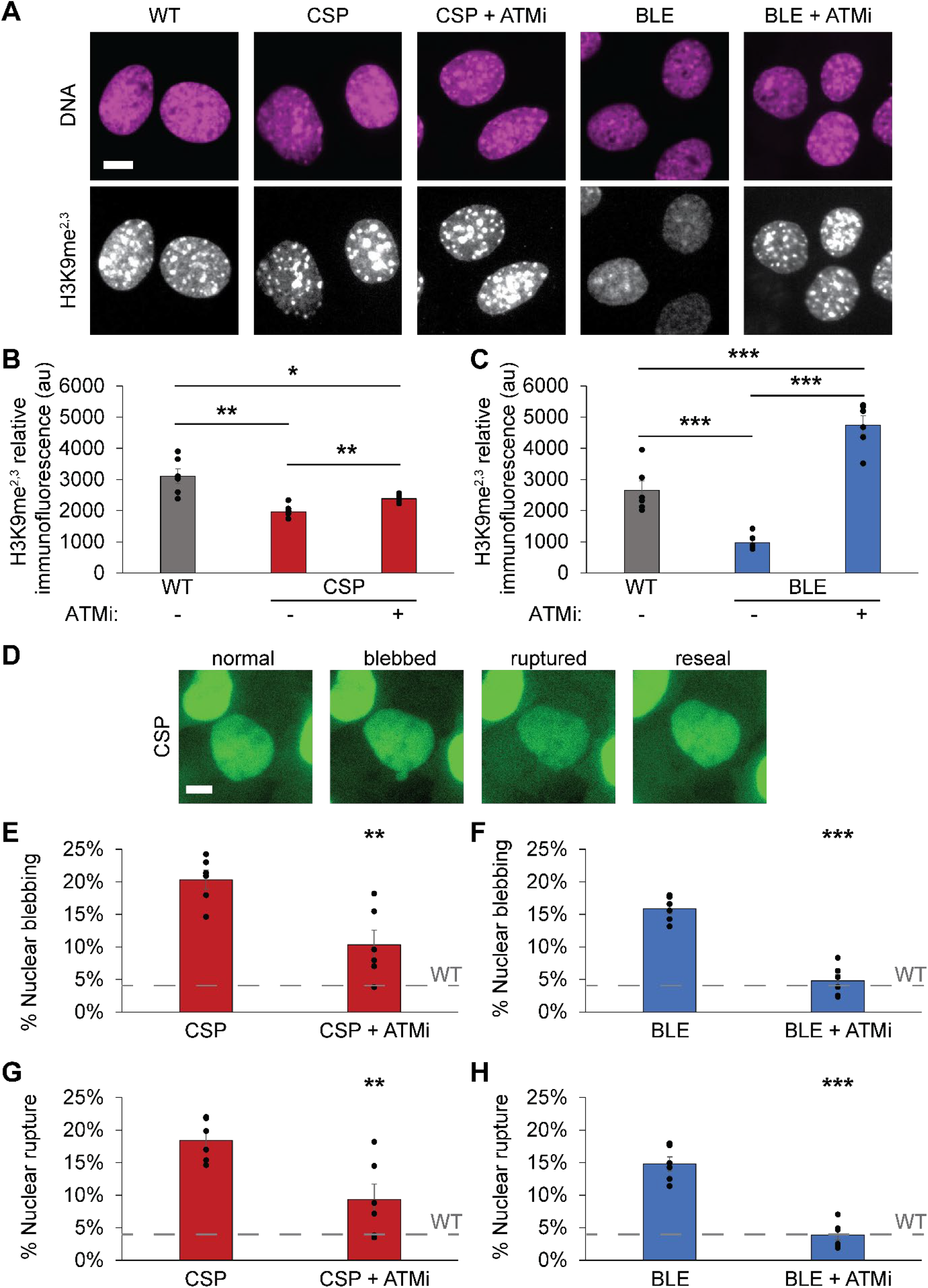
Induced DNA damage causes decreased heterochromatin that is responsible for nuclear blebbing and rupture. (A) Example images of nuclei stained with Hoechst (magenta) to label DNA and immunofluorescence for constitutive heterochromatin marker H3K9me^2,3^ (white) in wild type (WT), DNA damage treatment (CSP or BLE), and plus ATM inhibitor KU55933 (+ ATMi). ATMi was added one hour before DNA damage drug treatment for four hours. Graphs of (B) cisplatin high and (C) cisplatin compared to both wild type and with ATMi dual treatment in arbitrary fluorescent units (au). Each condition consists of 6 replicates with an average n > 45 nuclei each. (D) Example image of bleb formation and rupture in CSP high DNA damage drug treatment imaged via NLS-GFP. Graphs of nuclear blebbing percentages in (E) cisplatin high and (F) bleomycin without and with ATM inhibition by KU55933 (+ ATMi) for 16 hours. Graphs of nuclear rupture percentages in (E) cisplatin high and (F) bleomycin without and with ATM inhibition by KU55933 (+ ATMi). Dotted line denotes average wild type (WT) nuclear blebbing and ruptures around 4% to compare level of rescue upon ATMi. Each condition consists of 6 replicates n averages > 85 nuclei each for nuclear blebbing and rupture. Student’s t-test p values reported as *<0.05, **<0.01, ***<0.001, or ns denotes no significance, p>0.05. Error bars represent standard error. Scale bar = 10 µm.

Next, we tested the hypothesis that inhibition of ATM could rescue loss of heterochromatin and restore nuclear shape and integrity. Both nuclear blebbing and rupture caused by cisplatin or bleomycin were significantly decreased by dual treatment with ATMi (**Figure 3D-H**).

Interestingly, the small restoration of heterochromatin in cisplatin by ATMi led to a modest decrease in nuclear blebbing but was still more blebbing than in wild type (**Figure 3, E and G**). On the other hand, the drastic and greater than wild type increase in heterochromatin in bleomycin with ATMi treatment led to a more drastic rescue of nuclear blebbing and rupture returning them to wild type levels (**Figure 3, F and H**). These results suggest that DNA damage ATM-dependent signaling decreases heterochromatin to drive loss of nuclear rigidity, shape, and integrity.

There are well-established mechanism for changes in heterochromatin driven by ATM which is activated by DNA damage from both cisplatin (Colton *et al*., 2006) and bleomycin (Turenne *et al*., 2001). Both the magnitude of decreased H3K9me^2,3^ immunofluorescence and level of rescue provide unique insights. Interestingly, the magnitude of loss of heterochromatin upon DNA damage type does not appear to matter as both show an increased nuclear blebbing and rupture at similar levels (**Figure 1C-E** and **Figure 3D-H**). High cisplatin treatment, which largely causes DNA adducts, showed a modest decrease in heterochromatin and upon ATMi the rescue of heterochromatin was also modest. However, in DNA strand breakage via bleomycin treatment both the loss of heterochromatin and rescue upon ATMi dual treatment, returning to wild type or better levels, were more drastic than cisplatin. One possibility is that this difference in rescue could be due to the drastically higher level of DNA damage in cisplatin high compared to bleomycin (**Figure 1, A and B**). Alternatively, while bleomycin DNA strand breaks likely largely rely on ATM, cisplatin adducts could be activating another pathway, such as ATR, along with ATM to cause changes in nuclear physical properties. Overall, ATM-dependent response to DNA damage causes heterochromatin loss to result in a weaker nucleus that blebs and ruptures more frequently.

### DNA damage and nuclear integrity loss provide a negative feedback loop

Our finding that DNA damage affects nuclear mechanics, shape, and rupture reveals a new negative feedback loop (**Figure 4**), as it is already well-established that nuclear mechanics, shape, and rupture cause DNA damage (Stephens, 2020). The pathway follows that DNA damage leads to ATM-dependent decreases in heterochromatin (**Figure 3**) which results in chromatin-based nuclear softening (**Figure 2**) which ultimately results in nuclear blebbing, and rupture (**Figure 1**) established to lead to nuclear dysfunction. Specifically, nuclear rupture has been reported to cause DNA damage in non-migrating and non-artificially confined cells (Chen *et al*., 2018; Xia *et al*., 2018; Stephens *et al*., 2019b; Pho *et al*., 2023), migrating cells through confined spaces (Denais *et al*., 2016; Irianto *et al*., 2016; Raab *et al*., 2016; Pfeifer *et al*., 2018; Patteson *et al*., 2019), and externally compressed cells (Shah *et al*., 2021). Thus, more DNA damage to feedback into the outlined negative feedback loop (**Figure 4**). Other nuclear functions are only hypothesized to provide plausible negative feedback loops via transcription (Helfand *et al*., 2012; Berg *et al*., 2023) and cell cycle control (in preparation). However, we now provide evidence that DNA damage can cause nuclear blebbing and rupture which is established to lead to more nuclear dysfunction, including DNA damage. Thus, we postulate the nuclear blebbing and rupture provide negative feedback through DNA damage to drive a worsening disease state. This is only supported by the fact that nuclear blebbing increases correlate with disease severity in prostate cancer via the Gleason score (Helfand *et al*., 2012) while across the human disease spectrum loss of nuclear shape is a prognostic indicator (Gisselsson *et al*., 2001; Stephens *et al*., 2019a). Our report further details the interconnected relationship between nuclear function and physical properties via DNA damage response modulating nuclear mechanics, shape, and integrity which in turn affect nuclear functions.

**Figure 4.**
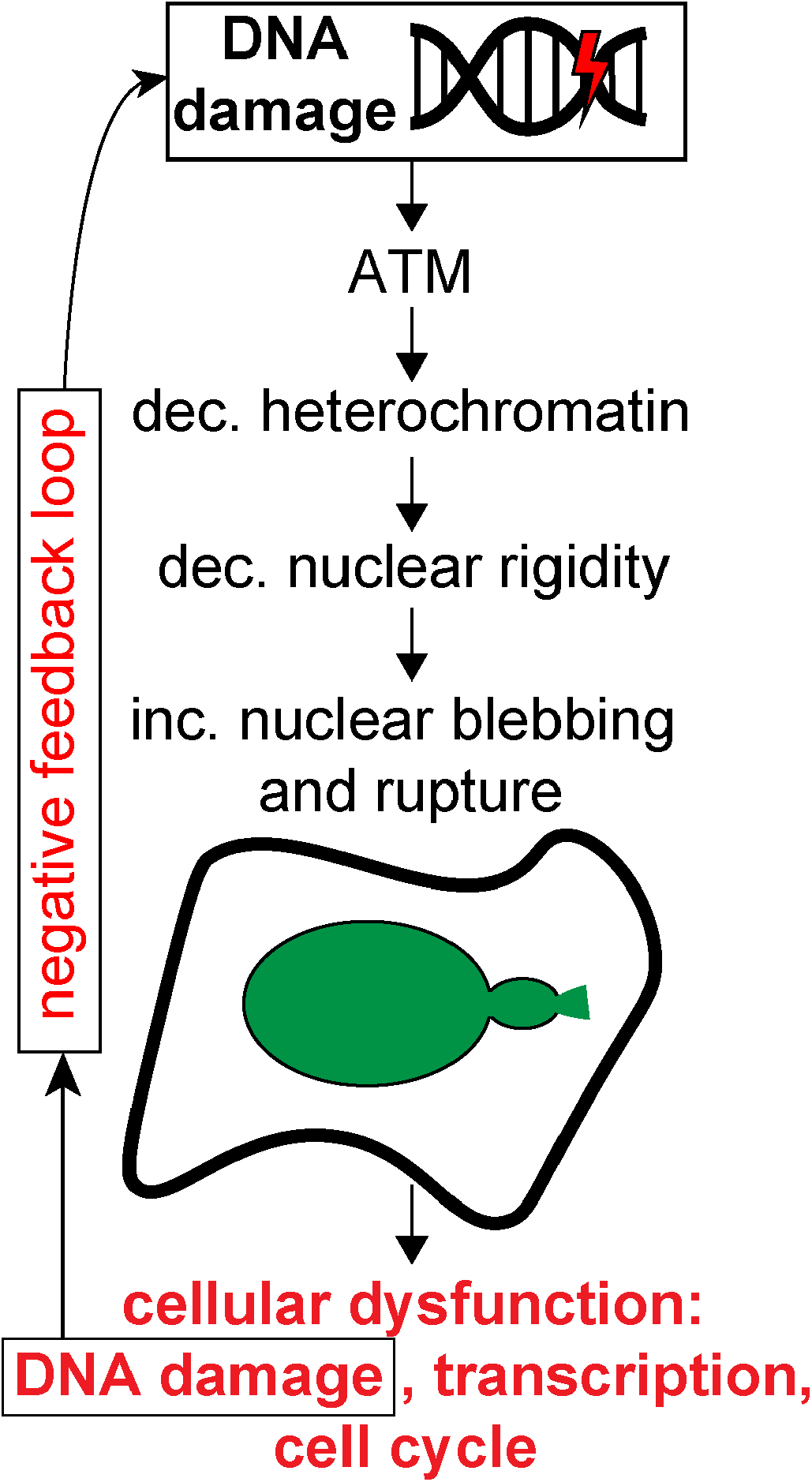
DNA damaged-based effects on nuclear mechanics, shape, and integrity provide negative feedback loop. The mechanism for DNA damage to alter nuclear physical properties that in turn causes nuclear rupture and cellular dysfunction that leads to negative feedback loop as it causes more DNA damage.

## Materials and Methods

### Cell culture

MEF WT NLS-GFP cells were cultured in a 60mm culture in DMEM (Corning) with 10% fetal bovine serum (FBS; HyClone) and 1% Penicillin/streptomycin (PS; Corning), incubated at 37°C and 5% CO2. Passaged after reaching 80-90% confluency. Cells were trypsinized, replated, and diluted with DMEM (Corning). HT1080 FUCCI human fibroblast cells (Marcus *et al*., 2015) were cultured similarly.

### Drug treatments

To induce DNA damage cells were treated with 5 µM or 25 µM of cisplatin (CSP; Tocris #2251) and 13 µM of bleomycin (BLE; Abcam ab142977) for 16 hours. ATM inhibitor KU55933 (abcam) was added at 20 µM for one hour before addition of DNA damage drug treatment.

### Time Lapse imaging and analysis

Images were acquired with Nikon Elements software on a Nikon Instruments Ti2-E microscope with Orca Fusion Gen III camera, Lumencor Aura III light engine, TMC CLeanBench air table, with 40× air objective (N.A 0.75, W.D. 0.66, MRH00401). Live cell time lapse imaging was possible using Nikon Perfect Focus System and Okolab heat, humidity, and CO_2_ stage top incubator (H301). Images with captured at 2-minute intervals for 16 hours using 30ms exposure time, 12-bit depth, and 3% FITCI (475 nm) power. Cells were treated with DNA damage drugs cisplatin 5 µM or 25 µM and bleomycin 13 µM 30 minutes before imaging. Time lapse images were analyzed by tracking bleb and rupture formation throughout the movie. To analyze, total number of blebbed nuclei, ruptured nuclei and number of nuclei per field of view were recorded. In order to calculate percent blebbed, and rupture, the number of blebbed nuclei were divided by the number of total nuclei per field of view. Then, average blebbed and rupture nuclei was used to compare differences between each treatment condition.

### Immunofluorescence

Cells were untreated or treated for 8 hours with biochemical and fixed with 4% paraformaldehyde in phosphate buffered saline for 15 minutes. Cells were permeabilized with 0.01% Triton X-100 (VWR Life Science) in PBS for 15 min, and washed with 0.006% Tween-20, (Promega) in PBS for 5 min followed by 3 washes with PBS. Cells were then blocked for one hour at room temperature with a blocking reagent 2% bovine serum albumin (BSA; Fisher Scientific) in PBS followed by staining with primary antibodies. Primary antibodies were diluted in the blocking solution at the following concentrations: H2K9ac 1:400 (9649, Cell Signaling Technologies), Lamin A/C 1:10,000 (4777, Cell Signaling Technologies), H3k9me^2-3^ 1:400 (5327, Cell Signaling Technologies), γH2AX-647 conjugated rabbit mAb 1:300 (9720, Cell Signaling Technologies), and γH2AX (9718, Cells were incubated with secondary antibodies Anti-mouse 555, Anti-rabbit 647, Anti-rabbit 555, Anti-rabbit Cy5 (Cell Signaling Technologies) in blocking solution for 1hr. Cells were then washed with PBS three times, 5 min each. Cells were stained with Hoechst 33342 (1:10,000, Invitrogen) in PBS for 15 min. Then, washed with PBS 3 times 5 min each.

### Fluorescence Imaging and analysis

Cells were imaged using Nikon Eclipse *Ti2* microscope, an objective lens Plan Abo Lambda 40x. Four excitation lights were selected, 405 nm DAPI with 5% power, 475 nm FITCI with 50% power, 555 nm TRITCI with 20% power, 635 nm Cy5 with 40% power. To analyze γH2AX data we followed previously established protocol by our lab (Pho *et al*., 2023). First, Hoechst DNA stain was used to determine nuclear ROIs. Then background was subtracted for the γH2AX channel. Next, using Nikon analysis software, Binary spot detection bright spots and bright clustered options were selected and a diameter of 0.5 µm was used. An empirically determined threshold was used across all conditions. The number of foci for each nucleus data was exported to an excel file. Statistical significance was determined using the t test.

To measure relative levels of euchromatin, heterochromatin, and lamin A/C, average intensities were acquired by measuring single nuclei as regions of interest after background subtraction from 30 x 30 pixel square areas containing no cells. For every biological replicate, measurements from 50-100 nuclei over were averaged to provide a single intensity measurement for each replicate experiment.

### Micromanipulation force measurements

As previously described (Stephens *et al*., 2017; Currey *et al*., 2022), vimentin null (V-/-) MEF nuclei were isolated from living cells with 0.05% Triton X-100 (VWR Life Science) in PBS. Using a pre-calibrated force micropipette and a pull micropipette, the isolated nucleus was grabbed, suspended, and the pull pipette was moved at 50 nm/s to provide a short 3 or long 6 µm extension of the nucleus. Measurement of the deflection of the force micropipette was multiplied by the bending modulus (1.2-2 nN/µm) to provide force measurement the nucleus as a nuclear spring constant (nN). The slopes of the force versus extension plots in the chromatin-dominated short regime (< 3 µm), chromatin-lamin domainted long regime (> 3 µm), and the difference between the nuclear spring constant of the short and long regime were plotted, and statistical significance was determined using the t test.

## Supporting information

Supplemental Tabel 1

## Acknowledgements

We would like to thank Pierre Vidi for helpful and insightful discussions and James Orth for sharing HT1080 FUCCI cell line. We also thank lab members Samantha Bunner, Kelsey Prince, and Katie Huang for their feedback and support. All are supported by the Pathway to Independence Award (R00GM123195) and Center for 3D Structure and Physics of the Genome 4DN2 grant (1UM1HG011536). The authors declare no competing interests.

