## Supplemental Figures 1 and 2 for "DNA damage causes ATM-dependent heterochromatin loss leading to nuclear softening, blebbing, and rupture"

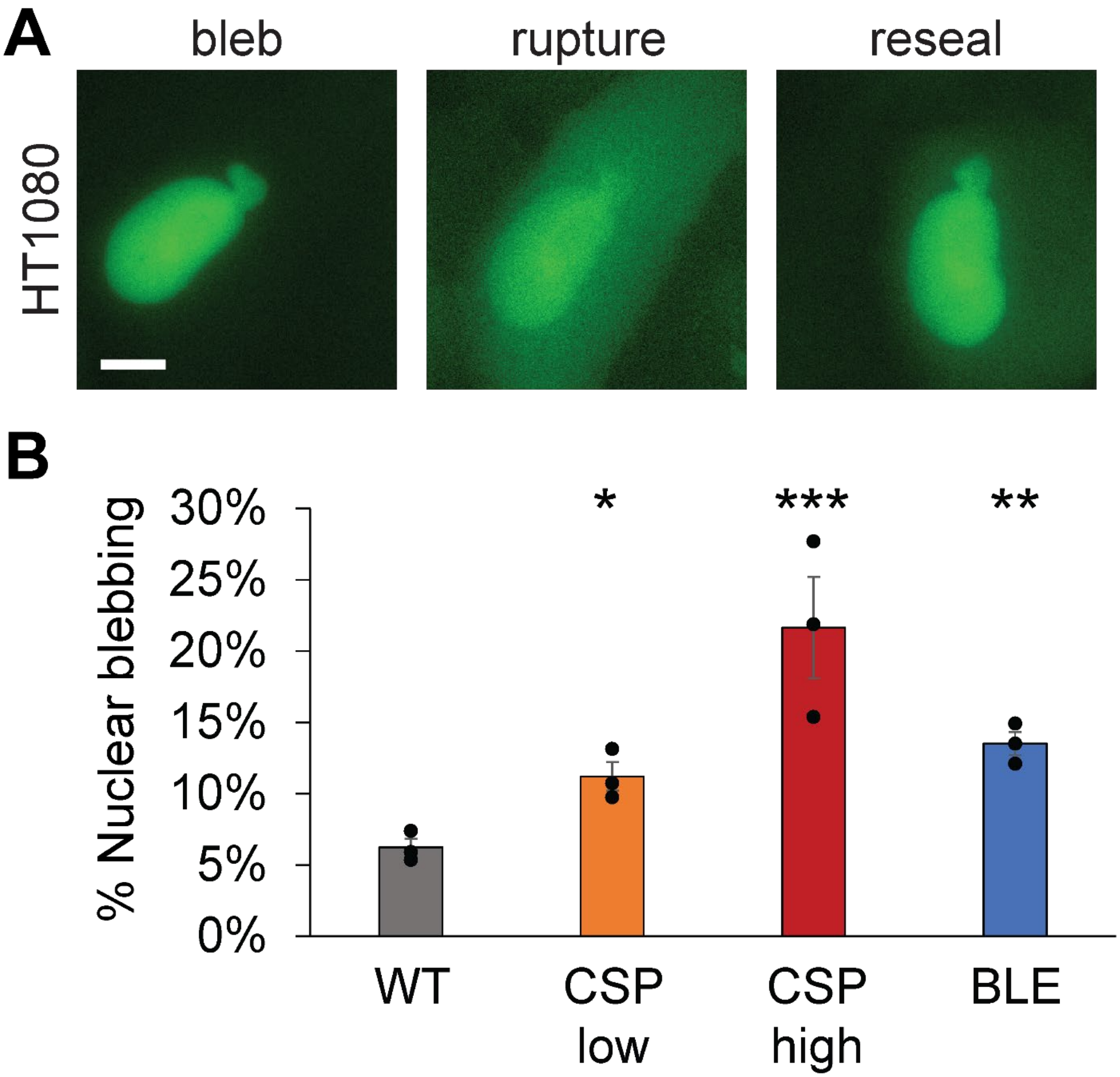


**Supplemental Figure 1. DNA damage induced by bleomycin affects nuclear blebbing more than cisplatin in HT1080 human cells.** (A) Example images of diffusible Geminin-GFP in HT1080 human cells showing nuclear blebs, rupture, and resealing. (B) Graph of nuclear blebbing for wild type (WT) and DNA damage treatments for 16 hours via cisplatin (CSP) and bleomycin (BLE).   Each condition consists of 3 replicates n = 30-100 cells each. Student's t-test p values reported as *<0.05, **<0.01, ***<0.001, or ns denotes no significance, p>0.05.. Error bars represent standard error. Scale bar = 10 µm.

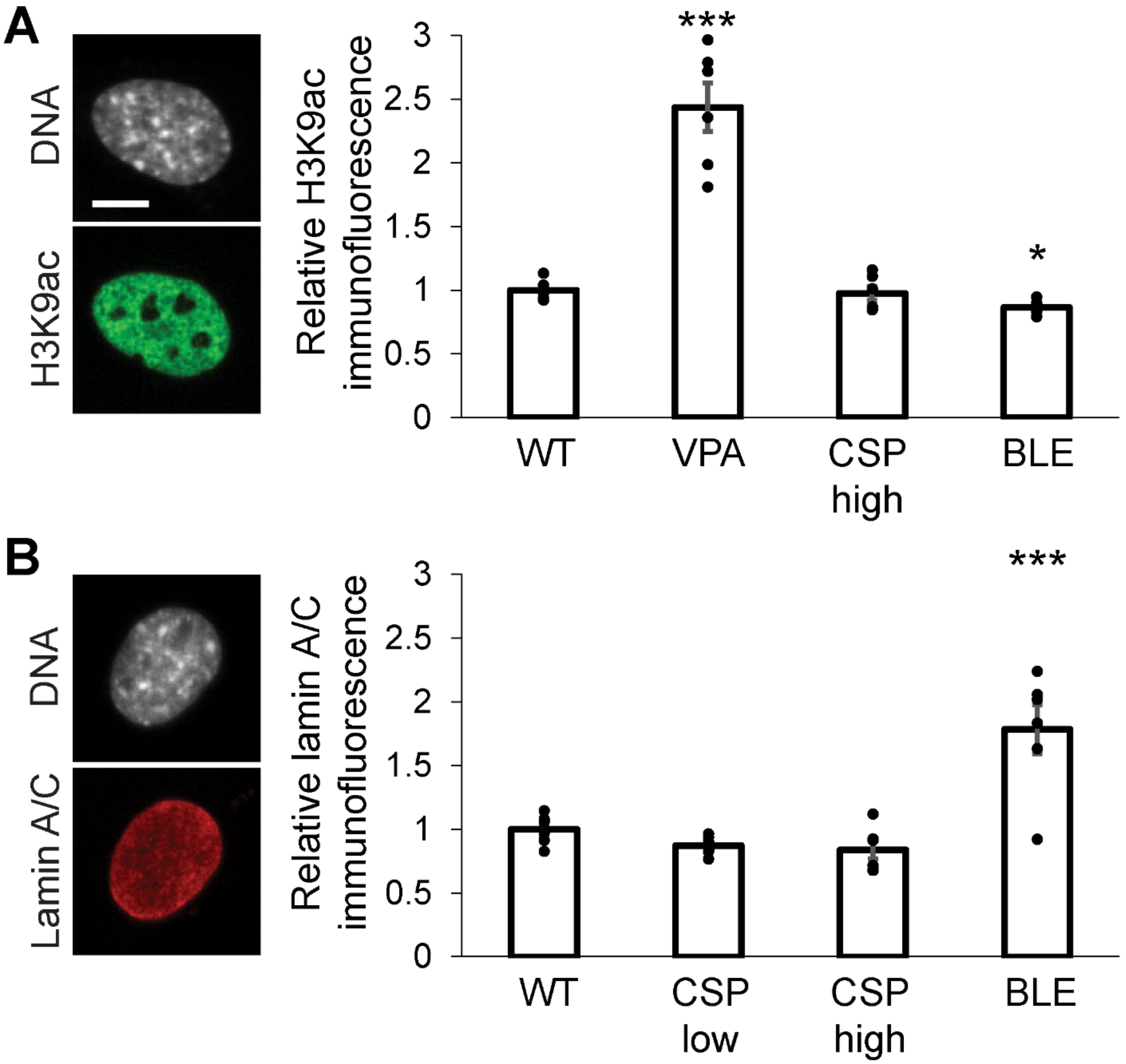


**Supplemental Figure 2. DNA damage drug treatments show no increase in euchromatin or loss of lamin A/C.** (A) Example image of wild type MEF nucleus DNA labeled via Hoechst and euchromatin labeled by immunofluorescence of H3K9ac. Graph of relative immunofluorescence of H3K9ac for control wild type (WT), increase via positive control via a histone deacetylase inhibitor (VPA), or DNA damage drugs cisplatin (CSP) and bleomycin (BLE). (B) Example image of wild type MEF nucleus DNA labeled via Hoechst and lamin A/C labeled via immunofluorescence. Graph of relative immunofluorescence of H3K9ac for control wild type (WT) and DNA damage drugs cisplatin (CSP) and bleomycin (BLE). Each condition consists of 6 replicates n > 20 cells each. Student's t-test p values reported as *<0.05, **<0.01, ***<0.001, or ns denotes no significance, p>0.05. Error bars represent standard error. Scale bar = 10 µm.

**Supplemental Table 1. Raw data.** This excel doc is a compiled document of all raw numbers used in the figures throughout the paper.
